# Transposable elements create distinct genomic niches for effector evolution among *Magnaporthe oryzae* lineages

**DOI:** 10.1101/2025.05.23.655716

**Authors:** Ana Margarida Sampaio, Daniel Croll

## Abstract

Plant-pathogen interactions are characterized by evolutionary arms races. At the molecular level, fungal effectors can target important plant functions, while plants evolve to improve effector recognition. Rapid evolution in genes encoding effectors can be facilitated by transposable elements (TEs). In *Magnaporthe oryzae*, the causal agent of blast disease in several cereals and grasses, TEs play important roles in chromosomal evolution as well as the gain or loss of effector genes in host specialized lineages. However, a global understanding of TE dynamics driving effector evolution at population scale and across lineages is lacking. Here, we focus on 16 *AVR* effector loci assessed across a global sampling of 11 reference genomes and 447 newly generated draft genome assemblies across all major *M. oryzae* lineages and outgroups. We classified each effector based on evidence for duplication, deletion and translocation processes among lineages. Next, we determined *AVR* gain and loss dynamics across lineages allowing for a broad categorization of effector dynamics. Each *AVR* was integrated in a distinct genomic niche determined by the TE activity profile contributing to the diversification at the locus. We quantified TE contributions to effector niches and found that TE identity helped diversify *AVR* loci. We used the large genomic dataset to recapitulate the evolution of the rice blast *AVR1-CO39* locus. Taken together, our work demonstrates how TE dynamics are an integral component of *M. oryzae* effector evolution, likely facilitating escape from host recognition. In-depth tracking of effector loci is a valuable tool to predict the durability of host resistance.

## Introduction

Plant-pathogen interactions are characterized by continuous evolutionary arms races, where hosts adapt to resist infection while pathogens evolve to overcome host defenses (Sironi et al., 2015). Effector-receptor dynamics are crucial for this process, where pathogens deploy effectors (*i.e.* avirulence factors, *AVR*) to manipulate host immunity (Lo Presti et al., 2015; Remick et al., 2023), while host receptors are responsible to detect them and trigger defense responses (Kanyuka & Rudd, 2019; Stukenbrock & McDonald, 2009). Genes encoding such effectors are among the most rapidly evolving genes in pathogen genomes, driven by high mutation rates and strong selection pressure (Sánchez-Vallet et al., 2018), enabling pathogens to evade host resistance and posing a challenge for effective disease control. Effector genes are often encoded in subtelomeric chromosomal regions, which are repeat-rich regions and evolve more rapidly than regions encoding housekeeping genes (Dong et al., 2015; Faino et al., 2016). These genomic regions are also frequently associated with transposable element (TE) activity, which can disrupt coding sequences or promote regulatory changes (Chuma et al., 2011; Fouché et al., 2018; Kang et al., 2001; Sampaio et al., 2025; Seidl & Thomma, 2017; Whisson et al., 2012). Moreover, these chromosomal regions are also exhibiting a higher propensity for stochastic epigenetic regulation (Aparicio et al., 1991; De Las Peñas et al., 2003; Fan et al., 2008), and chromosomal rearrangements resulting in rapid effector gene evolution (De Jonge et al., 2013; Shi-Kunne et al., 2018; Wang et al., 2020).

Plant pathogens capable of infecting both wild host plants and cultivated crops are of particular concern given their propensity to switch to new hosts (Anderson et al., 2004). *Magnaporthe oryzae* can infect over 50 wild and cultivated grass species including major cereal crops such as rice (*Oryza sativa*) and wheat (*Triticum aestivum*) (Wilson & Talbot, 2009). Despite the wide host range, *M. oryzae* genotypes are grouping into host-specialized forms and recognized as different pathotypes (Choi et al., 2013; Kim et al., 2019; Valent, 1990). The most studied pathotypes include *M. oryzae* Oryza (MoO), *M. oryzae* Triticum (MoT), and *M. oryzae* Lolium (MoL), causing blast disease on rice, wheat and ryegrass, respectively. The origin of new pathotypes was triggered by host jumps including the emergence of wheat and rice blast disease (Couch et al., 2005; Inoue et al., 2017). Wheat blast emerged from a host jump from ryegrass (*Lolium* spp.) to Brazilian wheat cultivars lacking the *RWT3* gene, which is the resistance gene expressing the receptor recognizing the PWT3 effector. While both *PWT3* and *PWT4* effector genes are found in the Lolium pathotype, cultivation of *rwt3* wheat cultivars (lacking the ability to recognize PWT3) allowed the emergence of a new Triticum pathotype isolates carrying the PWT3 effector but losing PWT4, an effector that would be recognized due to the presence of its complementary resistance gene (*RWT4*) in wheat cultivars. Subsequent loss-of-function mutations arose due to the nearby cultivation of wheat cultivars carrying *RWT3*, along with the spread of pathogens to common wheat varieties (Inoue et al., 2017). These findings support the hypothesis that host specialization was mainly driven by genetic changes at effector gene loci including gain or loss of effector functions (Yoshida et al., 2016).

Numerous *M. oryzae AVRs* were cloned and characterized for their interaction with plant resistance factors (De Wit et al., 2009). Among these, AVR-Pita interacts with the cognate Pi-ta resistance protein and shows homology to fungal zinc-dependent metalloproteases (Jia et al., 2000; Orbach et al., 2000). AVR-ACE1 is involved in the production of secondary metabolites and activates the rice resistance factor Pi33 (Böhnert et al., 2004; Collemare et al., 2008). AVR-Piz-t is recognized by Piz-t and suppresses pathogen-associated molecular pattern (PAMP)-triggered immunity (Li et al., 2009; Park et al., 2012). PWL effectors are primarily encoded by MoO and are rapidly evolving, small, glycine-rich secreted proteins (Kang et al., 1995; Sweigard et al., 1995). Beyond functional differences in their encoded proteins, *AVR* genes exhibit a high degree of genetic instability and are often localized in telomeric regions (Farman, 2007; Rehmeyer et al., 2006). Such high rates of sequence changes are likely increasing their adaptative potential to evade host recognition. Sequence diversification occurred mostly through simple point mutations as reported for *AVR-Pita* and *AVR-Pik* (Dai et al., 2010; Kanzaki et al., 2012; Longya et al., 2019), or sequence rearrangements causing segmental deletion of coding sequences as for *AVR-Pita* and *AVR-Pib* (Orbach et al., 2000; Zhang et al., 2015) associated with gains in virulence. There is also evidence for horizontal transfer of *PWT4* from *M. pennisetigena* to an Avena isolate from Brazil (Inoue et al., 2021). There is strong evidence that TEs impacted *AVR* loci and facilitate rearrangements. TEs facilitated virulence gains of MoO through loss-of-function mutations such as the insertion of a Mg-SINE into the *AvrPi9* coding sequence (Wu et al., 2015) or gains of virulence linked to *AVR-Pita* and *AVR-Pib* due to a Pot3 TE insertion (Hu et al., 2022; Kang et al., 2001). TEs were also likely facilitating the translocations observed for several *AVR* genes including *AVR-Pita* (Chuma et al., 2011). The loss of telomeric ends resulting in the elimination of *AVR-Pita* and *AVR-Pii* (Chuma et al., 2011; Khang et al., 2008) were likely also favored by the repetitive nature of subtelomeric regions. Overall, the rapid evolution of *M. oryzae AVRs* to evade host recognition is likely facilitated by TE dynamics. TEs have specifically expanded in MoO compared to MoT and MoL (Lin et al., 2024; Nakamoto et al., 2023). Furthermore, TE insertions also mediated the divergence of *M. oryzae* populations infecting different rice subspecies (Lin et al., 2024; Nakamoto et al., 2023). However, a population genomics perspective on TE impacts on effectors leveraging the vast available genomic datasets on *M. oryzae* lineages is lacking.

Here, we used extensive genomic datasets covering all major *M. oryzae* host-associated pathotypes to recapitulate insertion dynamics near 16 *AVRs* and included *M. grisea* and *M. pennisetigena* as outgroups. We recapitulated chromosomal rearrangements affecting *AVR* loci in the *M. oryzae* pathotypes MoO, MoT, and MoL using reference-quality genomes. We combined synteny analyses with draft assemblies from available sequencing datasets to assess *AVR* gain/loss patterns in conjunction with TE insertion dynamics surrounding the *AVRs*.

## Material and Methods

### *Magnaporthe* genomic datasets and genome assemblies

We performed analyses on a global collection of genomic datasets comprising 458 *Magnaporthe* isolates (Supplementary Table S1 and S2). All genomes were accessed from public databases reported by previous studies (Gladieux et al., 2018; Latorre et al., 2023; Thierry et al., 2022; Zhong et al., 2018). The geographic origin of isolates and the host organism information was retrieved from metadata attributes and cross-checked with the associated literature. The isolates were from Asia (n=256), South America (n=96), Africa (n=69), Europe (n=11), North America (n=8), and a small number without reported origin (n=18). Out of the 458 isolates, 448 belonged to *M. oryzae* collected from 18 different host species, including cereals and grasses. We also included as outgroups two isolates of *M. pennisetigena* (collected on *Pennisetum* sp.) and eight isolates of *M. grisea* isolated from *Cenchrus* and *Digitaria* sp. Overall, 447 isolates were sequenced using Illumina paired-end whole-genome sequencing (WGS) (Supplementary Table S1). Illumina sequencing data was initially filtered using fastp v0.23.4 with default settings (Chen et al., 2018) to remove adapter sequence and low-quality reads. *De novo* draft assemblies were generated using the software SPAdes v3.15.5 (Bankevich et al., 2012) with the “careful” method and automated k-mer selection. All genomes were verified to have more than 95% completeness using BUSCO version 5.8.2 (Simão et al., 2015) searching the Ascomycota orthology database. We used QUAST to calculate assembly metrics (Gurevich et al., 2013). We retained assemblies with N50 (length of the shortest contig for which longer length contigs cover at least 50% of the assembly) above 17,034 bp and a total assembly size above 36.72 Mbp. Eleven *Magnaporthe* reference-quality genome assemblies were also included in this study (Dean et al., 2005; Gomez-Luciano et al., 2019; Liu et al., 2024; Peng et al., 2019). These genomes included seven *M. oryzae* genomes (3 MoO, 3 MoT, 1 MoL) and four outgroup genomes (2 *M. grisea* and 2 *M. pennisetigena*) (Supplementary Table S2).

### Phylogenetic analyses

Phylogenetic relationships were assessed separately for the set of *Magnaporthe* reference-quality genomes and for 405 *Magnaporthe* draft genomes associated with the most frequently sampled host species. For the phylogenomic tree, we used single-copy genes predicted by AUGUSTUS v3.5.0 (Stanke et al., 2004) using the pretrained gene prediction database available for the *M. grisea* genome. Predicted protein sequences were used for orthology analyses performed with Orthofinder v2.5.5 (Emms & Kelly, 2019). Single-copy orthologs present in at least 90% of the total number of isolates were kept. For the phylogenetic reconstruction of the large *Magnaporthe* worldwide collection, we retained 300 randomly selected single-copy orthologs to reduce computational load. Selected ortholog protein sequences were aligned as a supermatrix using the AMAS tool v1.0 (Borowiec, 2016). Phylogenetic trees were built using RAxML v8 (Stamatakis, 2014) to construct a maximum-likelihood phylogenetic tree with the parameters -m PROTGAMMAAUTO for protein sequences with 1000 bootstrap replicates.

### Identification of effector homologues

We analyzed the 447 draft genomes produced by SPAdes and the 11 reference-quality genomes (including outgroups) to search for homologs of 16 *M. oryzae* effectors. We focused on cloned and well-characterized effectors identified in Oryza or Triticum pathotype isolates (Table 1). For effectors present in more than one pathotype, effector sequences with best hits in the reference-quality genomes were used as query for BLASTn analyses (Camacho et al., 2009). Hits were filtered for a maximum *e*-value of 10^-5^, followed by individual minimum length filtering for each effector based on visual inspection of alignment length distributions. The ORF finder tool (https://www.ncbi.nlm.nih.gov/orffinder/) was used to refine effector open reading frames detected by BLASTn hits.

**Table 1:**
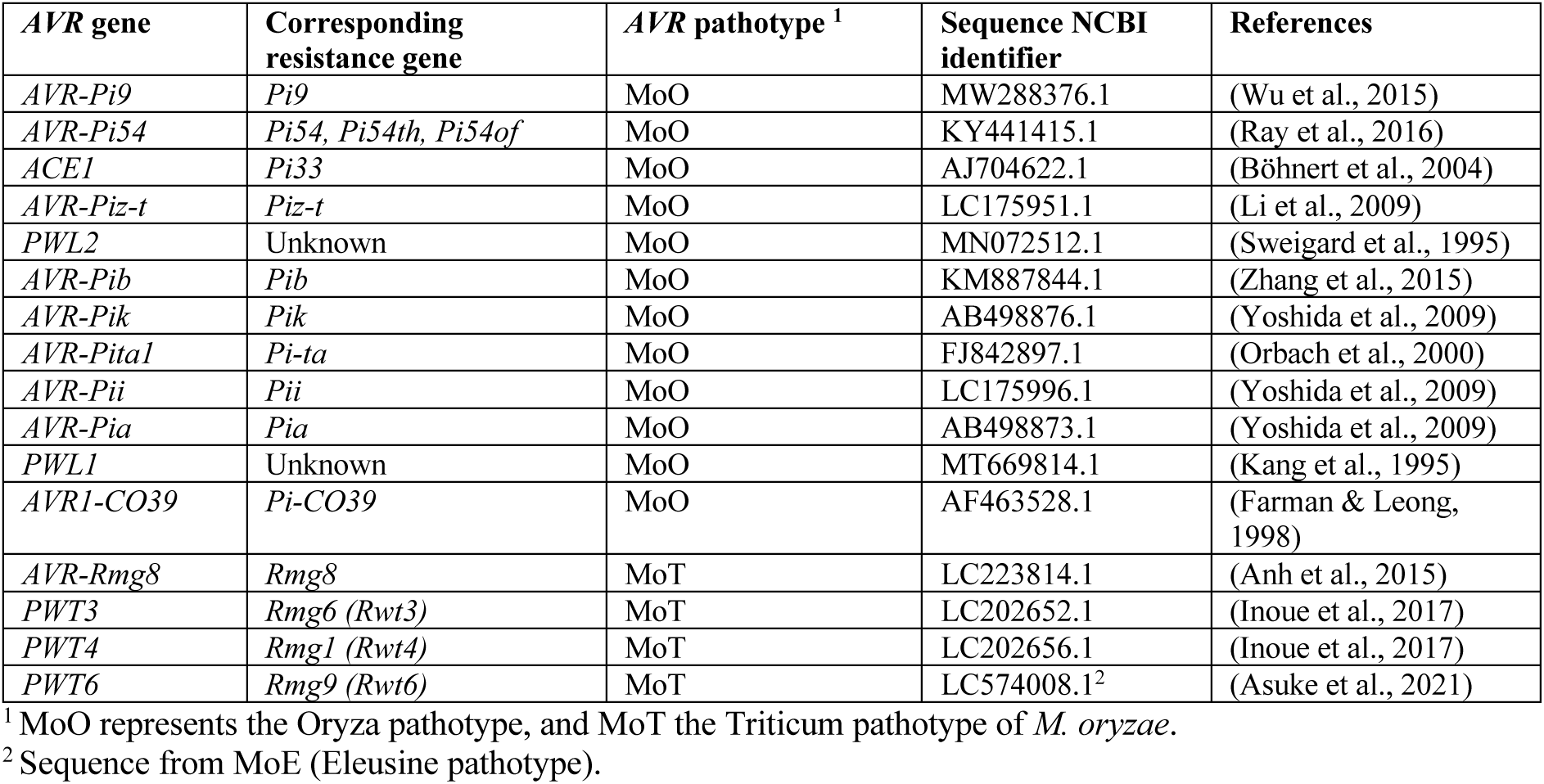
Overview of *M. oryzae AVR* effectors analyzed in this study. The *AVR* pathotype identifies the *M. oryzae* pathotype in which the effector was first characterized.

### Transposable element annotation and analysis

To detect TE insertions near effectors, all draft assemblies, reference-quality and outgroup genomes were annotated with the TE consensus sequences reported for *Magnaporthe* (Lin et al., 2024) (available from https://github.com/S-t-ing/mBio-data-availablility/blob/main/Mo.TE_Consensus.fasta). For this, we used RepeatMasker v4.0.9_p2 with “-no_is” and “-nolow” parameters to skip simple repeat and low complexity region annotations. Annotated TEs shorter than 50 bp were filtered out. The search for TEs near effector loci was restricted to a 1000 bp window up- and downstream the effector gene. Only MoO and MoT isolates clearly identified in large clades based on the phylogenetic tree were retained these analyses. Percentage of TE superfamilies per *AVR* in MoO and MoT was calculated per effector based on the total TE number per pathotype. Percentage of TE superfamilies per insertion site (bp) has been calculated per effector based on the total number of isolates containing at least one TE for this *AVR*. We used the ANOVA function in *R* to test for the impact of TE family, *AVR* identity, and the *AVR* pathotype (MoO and MoT) on variation in TE counts near effectors. The proportion of variance explained by each factor and interactions was calculated based on the sum of squares.

### Analyses of the *AVR1-CO39* locus

To assess the *AVR1-CO39* locus organization in genomes of different pathotypes, we first analyzed reference-quality genomes. Synteny was plotted with genoplotR v0.8.11 (Guy et al., 2010) for the *Magnaporthe* reference genomes using gene and TE annotation (Lin et al., 2024). For *M. oryzae* isolates, we mapped coding sequences from the MoO 70-15 (GCF_000002495.2) gene annotation (Dean et al., 2005), and for outgroup *M. pennisetigena* we mapped coding sequences from the Br36 (GCA_004337985.1) gene annotation (Gomez-Luciano et al., 2019). The two annotations of 70-15 and r36 provided complementary coverage of major haplotypes found at the locus. AUGUSTUS gene annotations as described above were used to infer coding sequences in non-*MoO* isolates. *Magnaporthe* genomes assembly having *AVR1-CO39* were inspected for TE presence surrounding the effector gene.

## Results

### Samples distribution, genome assembling and reference genomes

To unravel *M. oryzae AVR* evolution, we assembled a collection of genomic datasets for 458 *Magnaporthe* spp. isolates collected across continents and diverse hosts (Figure 1A) (Gladieux et al., 2018; Latorre et al., 2023; Thierry et al., 2022; Zhong et al., 2018). Most isolates were collected in Asia (59%) and predominantly infecting *O. sativa* (65%) (Figure 1A), reflecting the high incidence of rice blast disease in this region. Isolates infecting cereals such as *Triticum* sp. as well as grasses such as *Lolium* sp. and *Eleusine* sp. across continents were also included (Figure 1A). From the 458 analyzed isolates, 448 belonged to *M. oryzae* (Supplementary Table S1). Genomic data included 441 short read sequencing datasets and 7 reference-quality genomes from MoO, MoL and MoT (Supplementary Table S2) (Figure 1B). Ten outgroup genomes from two distinct *Magnaporthe* species (*M. grisea* and *M. pennisetigena*) were also included in the analysis of which four genomes were of reference quality (Supplementary Table S2). The selected outgroup *Magnaporthe* species *M. grisea* and *M. pennisetigena* clustered separately from each other and separated well from *M. oryzae* in the phylogenomic tree analysis (Figure 1B). To complement the available reference genomes, we either accessed or assembled draft genomes for 447 additional isolates. Assembly genome sizes ranged from 39.1 Mbp in isolates infecting *Leersia* spp. and 42.9 Mbp in isolates infecting *Cenchrus* sp. (Figure 1C). Assembled genomes for this study showed acceptable contiguity for the purpose of analyzing coding regions with the N50 (length of the shortest contig for which longer length contigs cover at least 50% of the assembly) averaging between 19,433 bp in isolates infecting *Pennisetum* sp. and 103,607 bp in isolates infecting *Setaria* sp. (Figure 1C). Assembly genome sizes were not meaningfully correlated with the assembly contiguity (*i.e.* N50), which supports the notion that the totality of the genome is reasonably well covered despite variation in assembly quality (Supplementary Figure S1). The reference-quality genomes showed BUSCO completeness scores ranging from 96.5% in PM1 (*M. pennisetigena*) to 97.9% in *M. oryzae* isolates infecting *Oryza* spp. (Figure 1D). All draft genome assemblies similarly exhibited >95% BUSCO completeness and are hence comparable to the reference genomes in terms of gene content. A small number of *Oryza*-infecting isolates showed slightly lower BUSCO completeness scores compared to the reference genomes, yet the assembly genome sizes were similar.

**Figure 1:**
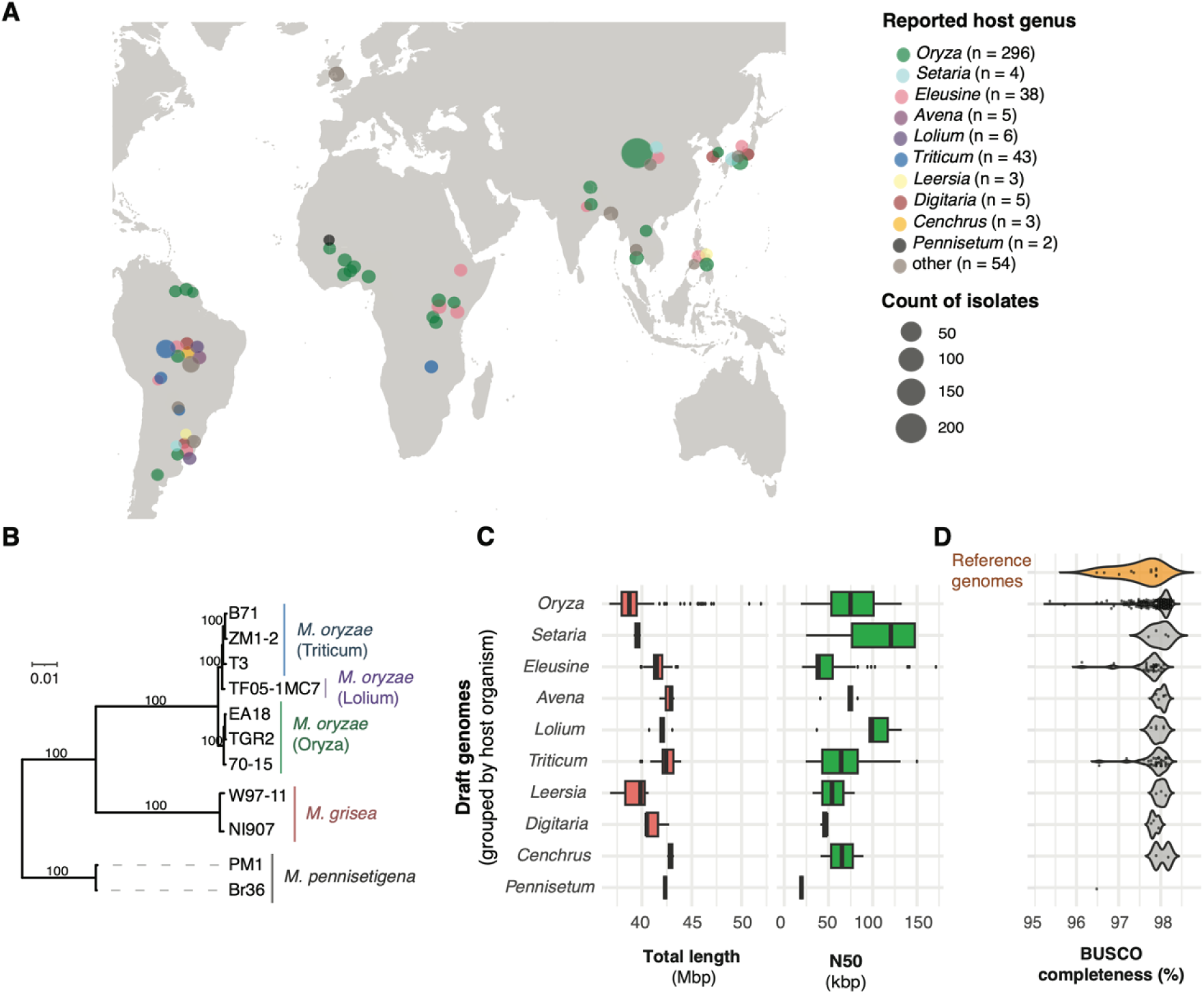
Global panel of analyzed *Magnaporthe* spp. genomes. A) Geographical distribution of the *Magnaporthe* spp. isolates. The color identifies the reported host genus, and the size defines the number of samples from the same location. B) Phylogenomic tree of reference-quality genomes. The host genus is reported in parentheses. Bootstrap confidence values >80% are displayed. C) *De novo* draft assembly genome size for genomes assembled from short-read sequencing data. Genomes are grouped by reported host genus. D) Analyses of genome completeness based on BUSCO completeness percentages for reference-quality genomes (orange on top) and *de novo* draft assembled genomes.

### Rearrangements of *AVRs* among *M. oryzae* reference genomes

Some *M. oryzae AVR* underwent chromosomal translocations. To comprehensively track *AVR* localization among isolates, we focused on 16 cloned and characterized *AVR* effectors (Table 1) in seven reference *M. oryzae* genomes from MoO, MoL and MoT. We expected MoO effectors to be shared among most MoO reference genomes. However, from the 12 MoO *AVR* effectors (Table 1), only seven were shared among all MoO reference genomes and only four were found in all *M. oryzae* reference genomes (Figure 2A). These four *AVRs* include *AVR-Pi9* and *AVR-Pi54* being at conserved chromosomal locations within pathotypes, while *ACE1* and *AVR-Pik* showed translocations among MoO isolates (Figure 2C). *AVR-Pita1*, *PWL1* and *PWL2* were present in all MoO reference genomes but have undergone duplication events (Figure 2D). We also identified deletions of two MoO *AVRs* (*AVR-Piz-t* and *AVR-Pib*) in at least one MoO reference genome (Figure 2E). *AVR-Pii* was absent in two MoO reference genomes (Figure 2E) but exhibited a duplication in two MoT reference genomes (Figure 2D). *AVR-Pii* was the only MoO effector that does not exhibit a duplication in a reference genome for its associated pathotype but in other *M. oryzae* pathotype reference genomes. *AVR1-CO39* and *AVR-Pia* suffered the most dramatic loss, being absent in all MoO reference genomes (Figure 2E). *AVR1-CO39* is shared among all MoL and MoT reference genomes though (Figure 2E). MoT effectors showed similar loss patterns to MoO effectors. Two MoT effectors were shared among the MoT reference genomes (Figure 2B). *PWT3* is shared among all *M. oryzae* reference genomes and located at a conserved position on chromosome 5 (Figure 2B). PWT4 is absent in most *M. oryzae* reference genomes except MoT. *PWT6* exhibited the most pronounced pattern of effector loss with the *AVR* being retained in only two MoO reference genomes (Figure 2D). Overall, *AVR* localizations are highly dynamic among the seven reference genomes with contributions by translocations, deletions and duplication events.

**Figure 2:**
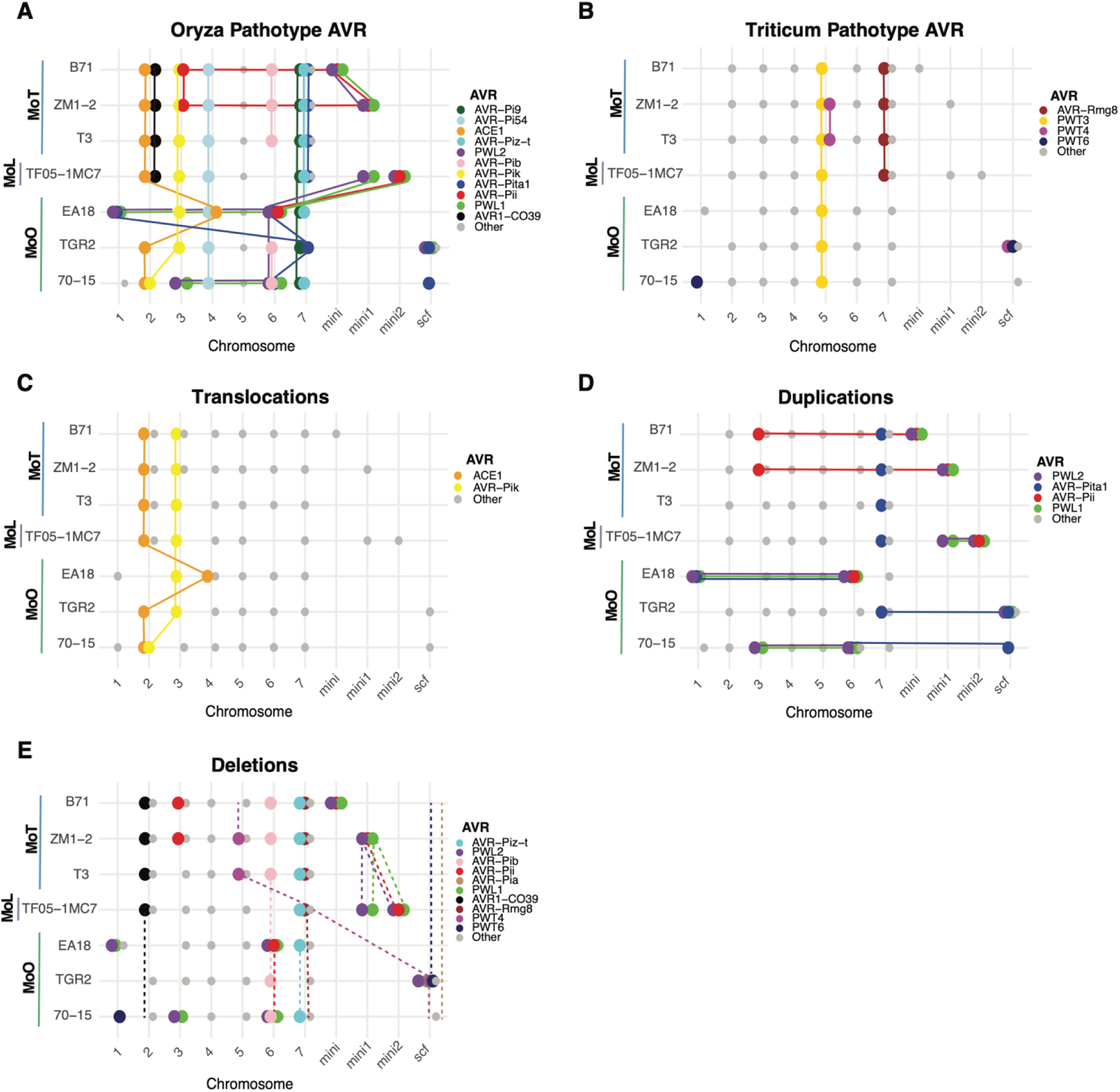
Overview of *AVR* effector synteny among *M. oryzae* reference genomes from MoO, MoL and MoT. Chromosomal localizations of *AVR*s are reported for (A) MoO and (B) MoT. Overview of *AVRs* showing evidence for (C) translocation among chromosomes, as well as (D) duplication and (E) deletion events. Dashed lines connect *AVR* localizations between reference genomes. If an effector was missing from a genome, a dashed line without a circle symbolizes absence of the effector gene.

### Effector gain and loss dynamics across global *M. oryzae* lineages

To comprehensively map the evolution trajectory of *AVRs* across *M. oryzae* lineages, we searched for effector homologs across a dataset of 458 *M. oryzae* genomes (Supplementary Table S3). Isolates infecting *Cenchrus* and *Pennisetum* sp. were used as outgroups to clarify gains and loss patterns. Relationships among isolates were assessed based on a phylogenomic tree including 300 single-copy orthologs. The phylogenetic grouping is consistent with previous studies of *M. oryzae* lineage diversification (Figure 3). Isolates belonging to the *Avena*, *Lolium* and *Triticum* pathotypes clustered together. Among those, isolates collected from *Triticum* sp. were the most dispersed across the tree, corroborating the high genetic diversity reported for MoT pathotype isolates (Rahnama et al., 2023). We verified consistency of phylogenetic placements for 24 isolates overlapping with a previous phylogenomic analysis (Gladieux et al., 2018). Three isolates reported as collected on *Oryza* sp. leaves (A-PHL-64, ARG-60 and ARG-61) (Onaga et al., 2020) did not cluster with the remaining MoO isolates. Furthermore, four MoT isolates were not clustering as closely with MoL and MoA as expected but rather with MoS (Figure 3).

**Figure 3:**
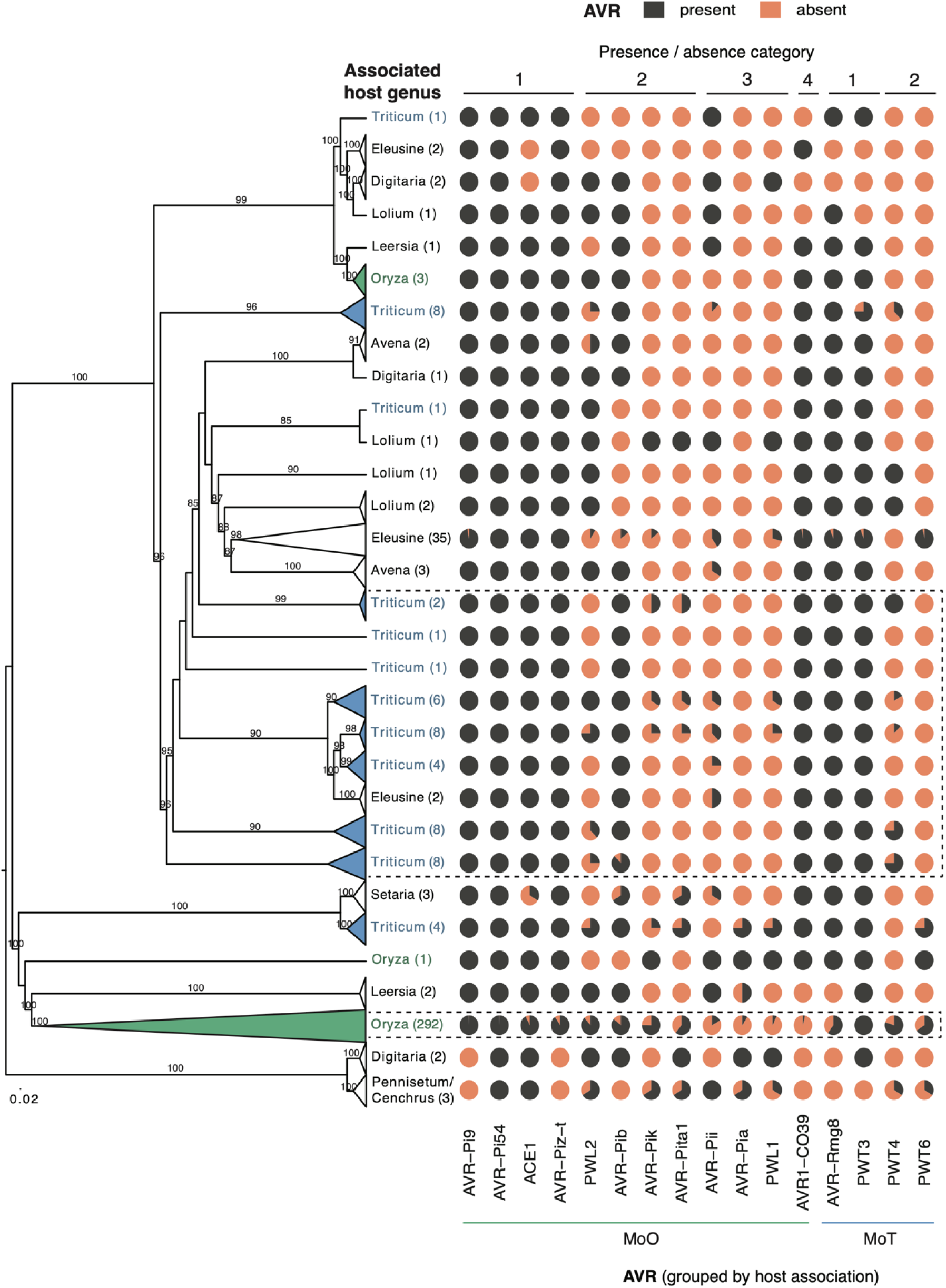
Effectors gain and loss dynamics across *Magnaporthe* spp. Maximum likelihood phylogenomic tree of 405 *Magnaporthe* genomes based on 300 protein sequence alignments of single-copy genes. Bootstrap confidence values >80% are displayed. Numbers in parentheses indicate the number of genomes. Pie charts indicate the proportion of isolates carrying specific effectors per clade. The presence/absence category refers to a summary classification of *AVR* gain/loss patterns observed across the phylogeny: 1) stable *AVR* presence; 2) *AVR* frequency increase in pathotype originally described for the *AVR* function; 3) general decrease in *AVR* frequency across *M. oryzae* pathotypes; 4) *AVR* frequency decrease in pathotype originally described for the *AVR* function.

To categorize effector gain and loss dynamics, we assessed the frequency of effectors according to their reported host (Table 1). Most *AVRs* were present in at least one of the outgroups infecting *Cenchrus* and *Pennisetum* sp., respectively, suggesting that most *M. oryzae* effectors were present in the common ancestor to all extant *M. oryzae* lineages. Exceptions include *AVR-Pi9*, *AVR-Piz-t*, *AVR1-CO39* and *AVR-Rmg8* being absent in the outgroups. We categorized *AVRs* into four types according to their frequency across *M. oryzae* lineages: 1) *AVRs* with largely stable frequencies among lineages; 2) *AVRs* increasing in frequency in the pathotype reportedly linked to the *AVR* function; 3) *AVR* largely lost in most *M. oryzae* pathotypes; 4) *AVR* largely lost in the pathotype reportedly linked to the *AVR* function. Among MoO isolates, all four gain/loss categories were observed for MoO effectors. *AVR-Pi9*, *AVR-Pi54*, *ACE1* and *AVR-Piz-t* are well conserved across the *M. oryzae* phylogeny (category 1; Figure 3). The near fixation of the *AVRs* suggests that recognition by the host is not widely distributed among host varieties or that the *AVR* serves an additional function.

### Transposable element colonization near *M. oryzae* effectors

The high frequency of rearrangements at *AVR* loci is consistent with high repetitive DNA content nearby. TE insertions are known to facilitate the creation of structural variation. Hence, we investigated patterns of TE dynamics surrounding *AVRs* and potential links to effector presence/absence variation. We annotated assembled contigs encoding the different *AVRs* for the presence of TE sequences considering a window of +/- 1000 bp of the effector coding sequence (Supplementary Table S4). The most frequent and conserved effectors (*AVR-Pi9*, *AVR-Pi54*, *ACE1* and *AVR-Piz-t*) loci were devoid or nearly devoid of TEs in proximity in both MoO and MoT isolates (Figure 4A and B). This suggests that the conservation of *AVR* effectors is facilitated by suppressed TE activity nearby. On the contrary, *AVR1-CO39*, an effector lost nearly entirely in MoO but remaining at high frequency in MoT isolates, showed the highest percentage of TEs among MoT isolates (Figure 4C). Hence, the high rate of TE insertions nearby could have facilitated the loss of the effector. Interestingly, the frequency of TE families is associated with the *AVR* identity. The two DNA transposons *POT2* and *POT3* are at high frequency near *PWT2*, *AVR-Pib* and *AVR-Pik* loci (Figure 4C). Retrotransposons were also abundant near specific *AVR*s. *LTR-RETRO5* and *LTR-RETRO7* were commonly found near the MoO effectors *AVR-Pita1*, *AVR-Pii*, and *AVR1-CO39*, while *LTR-RETRO6* was more frequent near the MoT effector *PWT6*. The *LTR-MGL3* retrotransposon showed a high frequency near *AVR1-CO39* and *PWT4*. Notably, *LTR-Pyret* was exclusively detected near *PWT3* in both MoO and MoT isolates. Additionally, the *MGL-LTR* retrotransposon was most frequent near effectors first characterized in MoT (Figure 4C). The distance between TEs and the nearby effector gene varied. Effectors in MoO had typically a higher distance to TEs than effectors in MoT isolates suggesting that sequence rearrangements could have affected the spacing between TEs and effectors (Figure 4A and B).

**Figure 4:**
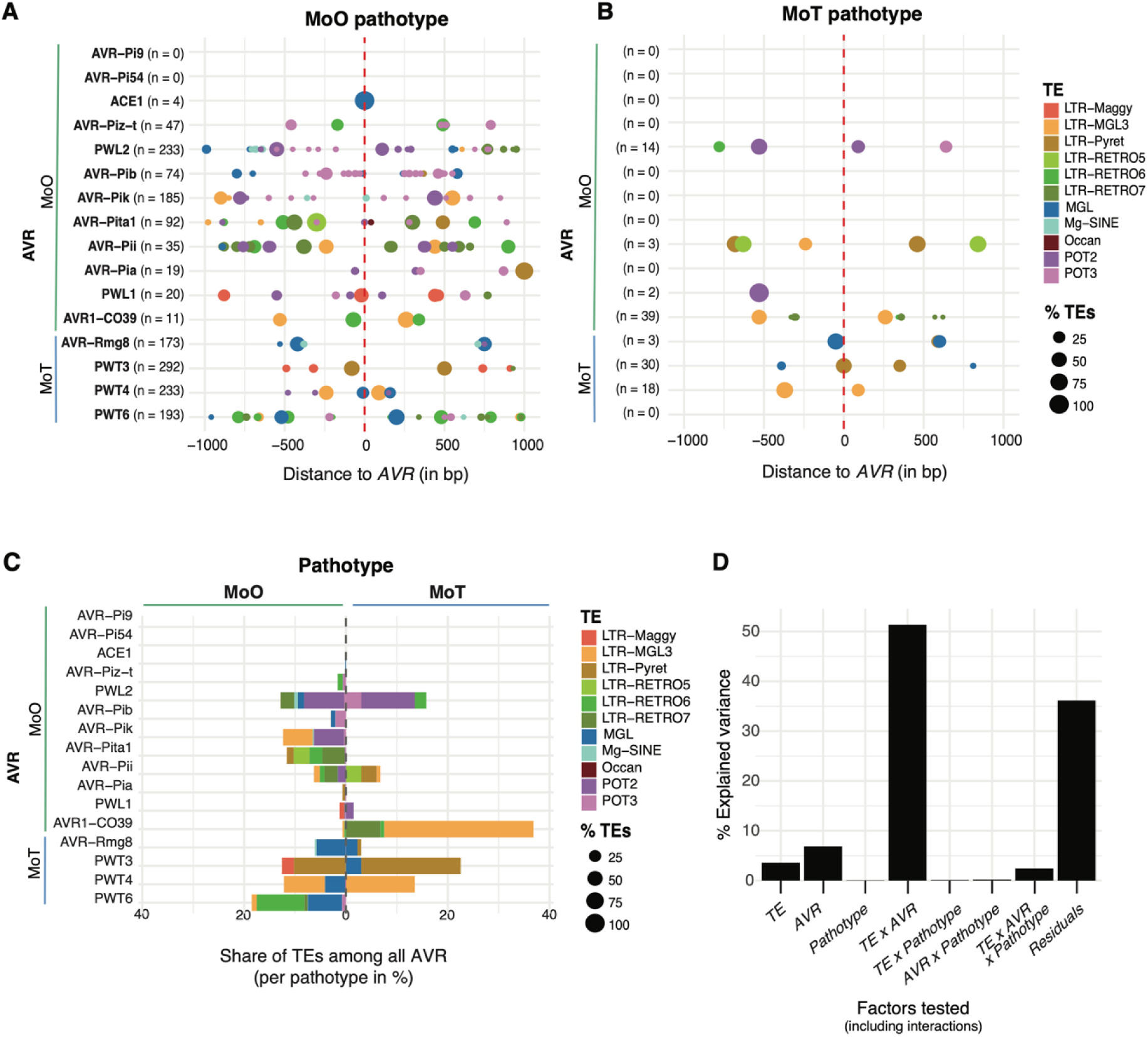
Transposable element (TE) dynamics and diversification near *M. oryzae AVR*. A) Presence of TEs in a +/- 1000 bp window surrounding *AVR* loci in Oryza pathotype (MoO) isolates. The circle color reflects TE identity and classification. The circle size indicates the percentage of isolates carrying the respective *AVR* having a specific TE present. 100% refers to all isolates with the *AVR* sharing a specific TE at a specific position. Counts (*n*) indicate the number of isolates exhibiting at least one TE for a given *AVR*. Negative bp distance values represent *AVR* upstream regions. B) Triticum pathotype (MoT) isolates. C) Relative abundance of TEs among *AVR* loci shown separately for MoO and MoT isolates. D) Analyses of variance (ANOVA) of factors potentially explaining variation in TE abundance among *AVRs*. The percentage of variance explained by the different models based on factorial combinations for TE, *AVR* and pathotype (for which the *AVR* was first described) and their interactions.

Given the heterogeneity in TE occupancy near *AVRs* across pathotypes, we sought to formally assess what factors explain best the variability in TE content. Using a multi-factorial ANOVA, we found that the TE identity (*i.e.* classification), the identity of the *AVR* and the pathotype explained a significant portion of the variance in TE occupancy (Figure 4D, Supplementary Table S5). The largest proportion of variance (51%) in TE occupancy was explained by the interaction of *AVR* and TE identity. This suggests that *AVR* loci may have co-evolved with distinct TE families, reinforcing the idea that certain TEs have affinity for specific chromosomal regions.

### *AVR1-CO39* locus dynamics

*AVR1-CO39* was the only effector at low frequency in isolates of the pathotype (MoO) in which the effector function was originally described and at high frequency in all the other pathotypes (Figure 3). *AVR1-CO39* was previously characterized for a sequence rearrangement at the origin of the host switch to rice (Farman et al., 2002; Peyyala & Farman, 2006; Tosa et al., 2005). The two variants were described as the G- and J-type in MoO lacking a functional *AVR* and an alternative W-type associated with an intact *AVR1-CO39* in isolates infecting weeping love grass (Farman et al., 2002). The most frequent type in MoO (G-type) is characterized by a complete loss of the coding sequence through a deletion and TE replacement. The J-type consists of a loss-of-function version caused by a repetitive element called REP1 (Farman et al., 2002). REP1 corresponds to LTR-RETRO6 in more recent TE annotations (Figure 5A). We analyzed the evolution of the *AVR1-CO39* locus organization across pathotypes by inspecting first the reference genomes. We used gene models annotated in the MoO genome 70-15 to obtain gene annotations in the additional *M. oryzae* reference genomes included in the comparison (Figure 5B). The W-type was represented by the MoL and MoT reference genomes (Figure 5B). The W-type carries genes adjacent to *AVR1-CO39*, which are absent in 70-15 (empty chromosomal region of the W-type; Figure 5B). We identified no locus synteny with the outgroup reference genome (Br36) (Figure 5B). As previously reported for the MoO isolate Guy11 (Farman et al., 2002), the 70-15 MoO reference genome shows a substantial contraction of the region adjacent to *AVR1-CO39* compared to the MoL and MoT genomes (Figure 5B).

**Figure 5:**
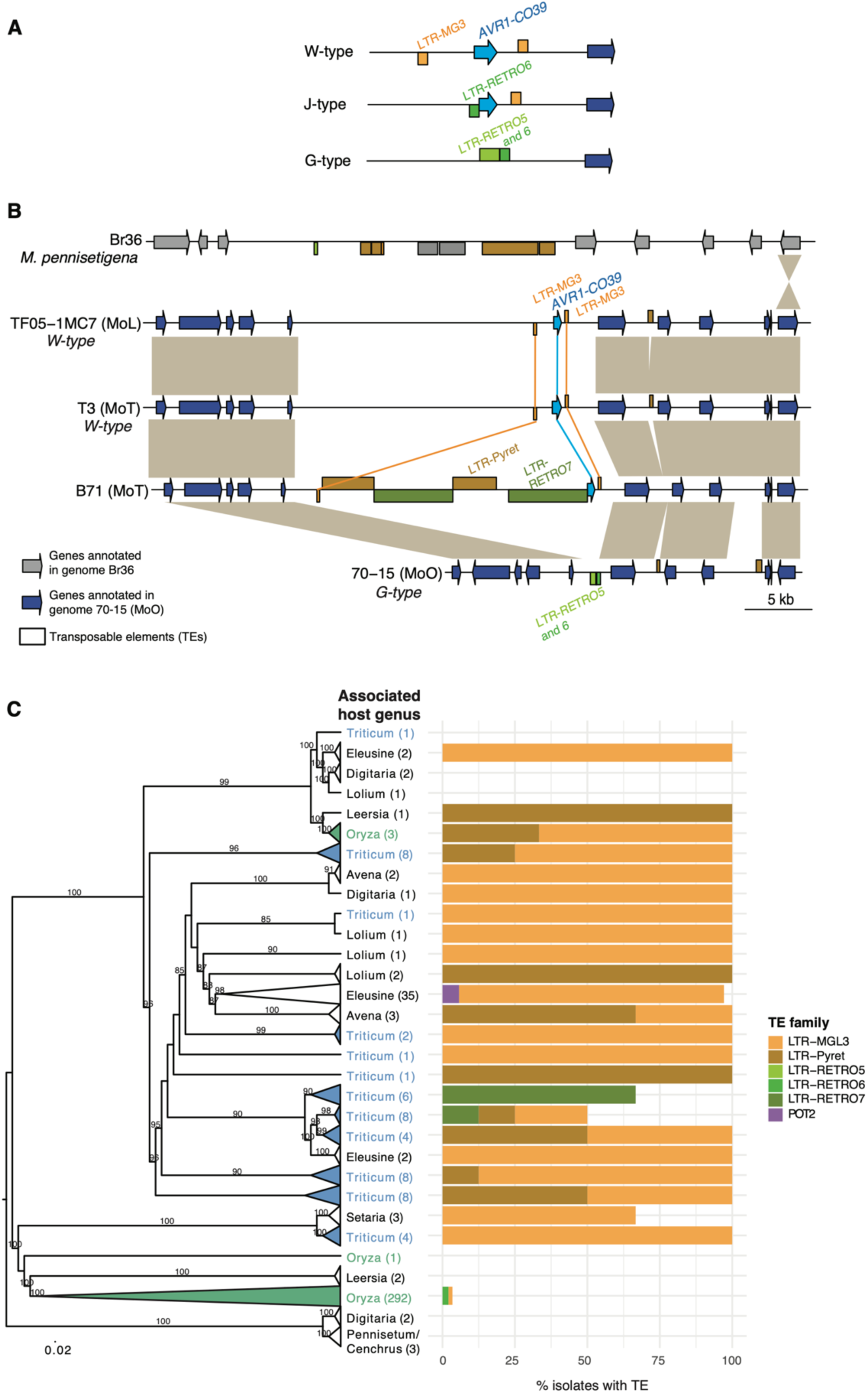
Chromosomal synteny analyses of the *AVR1-CO39* locus. A) Sketch of *AVR1-CO39* locus types as reported in the literature (Farman et al., 2002). B) Synteny plot of the *AVR1-CO39* region on chromosome 2 of MoO, MoL and MoT reference genomes and *M. pennisetigena*. Grey blocks between chromosomes indicate homologous regions. C) Comparison of TE types upstream of *AVR1-CO39* among *Magnaporthe* lineages. The phylogenomic tree was based on 300 single-copy orthologs. Bootstrap confidence values >80% are displayed.

The contracted region in 70-15 exhibits different TEs compared to the W*-*type, with *LTR-RETRO5* and *LTR-RETRO6* occupying the contracted region. The *LTR-RETRO5* has been previously identified in the G-type and the adjacent element was called *REP1*, which we identified here as *LTR-RETRO6*. The MoT isolate B71 exhibits yet distinct pattern with the *AVR1-CO39* upstream region fully occupied by TEs (Figure 5B). Two of these TEs were intact *LTR-RETRO7*, suggesting a recent insertion. One of the *LTR-RETRO7* insertion resulted in the truncation of *AVR1-CO39* in B71, differing in size from the J-type rearrangement. However, both retained ORF3, which encodes the *AVR1-CO39* domain interacting with the corresponding resistance gene (Ribot et al., 2013). The Zambia wheat blast outbreak isolate ZM1-2 (MoT) was reported to share a similar *AVR1-CO39* locus structure as B71 (Latorre et al., 2023). This highlights that the *AVR1-CO39* loss-of-function is not restricted to MoO isolates but is also occurring independently in MoT isolates through a different TE insertion. Since the different *AVR1-CO39* types are characterized by distinct TE content upstream of the coding sequence, we expanded our investigations to all isolates of the collection carrying the effector. We find that the *LTR-MGL3*, characteristic of the W-type rearrangement, was the most frequent among *Magnaporthe* lineages (Figure 5C). In contrast, the J-type was rare and found in only 10 MoO isolates (Figure 5C). The *LTR-RETRO7* characteristic of the J-type was found in the reference genome B71 and a small number of additional MoT isolates (Figure 5C). We identified two further TEs, *LTR-Pyret* and *POT2*, which were also found directly upstream of *AVR1-CO39* in our dataset. Our investigations of the locus indicate that the W-type is likely ancestral and has undergone multiple rearrangements followed by different TE insertions leading ultimately to the loss of the effector (*i.e.* G-type).

## Discussion

*AVR* effectors of *M. oryzae* have undergone rapid evolution to circumvent the matching plant receptors and increase pathogen virulence (Huang et al., 2014). Point mutations, insertions and deletions are widely described mechanisms responsible for the loss of avirulence function of *AVR* genes (Chuma et al., 2011; Hu et al., 2022; Orbach et al., 2000; Zhang et al., 2015). Abundance of TE sequences near effector regions can act as a mutagen or epigenetic regulator for accelerated effector evolution (Abraham et al., 2024; Bao et al., 2017; Jeon et al., 2015). In *M. oryzae*, TE content correlates with host identity and contributes to genetic differentiation among lineages (Lin et al., 2024; Nakamoto et al., 2023). Here, we show that effectors evolved well differentiated presence/absence patterns across *Magnaporthe* pathotypes, which at least in part reflects the diversity in recognition mechanisms by the different hosts. Across pathotypes of *M. oryzae*, we show that TE insertion dynamics most likely underpin the observed effector rearrangements. In contrast, conserved *AVRs* show no recent TE activity in the surrounding regions.

Variable *AVR* frequencies among *M. oryzae* host-specific pathotypes may be driven by adaptation to new hosts or the deployment of a new resistance gene. Here, we show that multiple MoO effectors were lost in different *M. oryzae* pathotypes including rice blast. This is consistent with the high rates of gene losses reported for rice-infecting *M. oryzae* compared to *Triticum* or *Avena* sp. infecting isolates (Yoshida et al., 2016). How closely the *AVR* frequencies among hosts reflects pathogenicity and recognition capabilities remains to be determined. Effector loss may be sufficient to escape host recognition and therefore contributing to host range expansion. This has been reported for instance for *PWT3*, with its loss coinciding with the widespread deployment of the complementary *R* gene *RWT3* (Inoue et al., 2017). In contrast, the more recently described *MAX* effectors do not necessarily reflect host specificity (Naour-Vernet et al., 2023). This is consistent with the absence of certain effectors, which may not reflect avoidance of host recognition. Well-conserved effectors across multiple pathotypes and an origin outside of *M. oryzae* may reflect a conserved function in virulence maintained by purifying selection. However, contributions to virulence may still vary among pathotypes, as observed for the *ACE1* effector (Collemare et al., 2008; Vy et al., 2024).

TEs represent approximately 10% of the *M. oryzae* genome (Bao et al., 2017; Dean et al., 2005) and can be inserted in or around *AVR* genes altering their virulence spectrum through transcriptional silencing, loss-of-function or loss of avirulence (Hu et al., 2022; Kang et al., 2001; Li et al., 2023; Li et al., 2009; Miki et al., 2009; Zhang et al., 2015). TE-mediated disruptions can result in the permanent loss of *AVR* genes, being a possible explanation for the observed deletion patterns. In *AVR-Pib*, a POT3 transposon insertion mediates effector loss-of-function in Philippine MoO isolates (Olukayode et al., 2019). We found a striking association of the TE *POT3* in the neighboring regions of *AVR-Pib* in MoO, suggesting that the TE plays a functional role. TE insertions can also promote the emergence of new virulent effector variants, as observed for the MGL retrotransposon insertion into the *ACE1* gene (Fudal et al., 2005) and POT3 insertion in *AVR-Pita1* (Kang et al., 2001; Zhou et al., 2007) and *AVR-Piz-t* (Li et al., 2009) ensuring the maintenance of such effectors. In our collection, all the four MoO isolates exhibiting TEs in the *ACE1* surrounding regions exhibited this same TE overlapping with the *ACE1* coding sequence. The remaining 270 MoO isolates were devoid of TEs near the effector. TEs can also facilitate the translocation of *AVR* genes such as *AVR-Pita* (Chuma et al., 2011). Here, we focused on MoO reference-quality genomes to investigate translocations of *ACE1* and *AVR-Pik*. However, we found no association between these translocations and specific TEs.

*M. oryzae* shows lineage-specific TE activity with LTR-retrotransposons expanded in certain lineages, rather than undergoing a single expansion followed by selective deletion (Nakamoto et al., 2023). TEs were often found near genes with presence/absence variation including effectors (Joubert & Krasileva, 2024). Here, we observed that well maintained *AVRs* had almost no TE activity in the surrounding area. This is possibly explained by their chromosomal location. In contrast to most of the *M. oryzae AVR* genes, usually located in telomeric or subtelomeric regions (Chen et al., 2007; Chuma et al., 2010, 2011; Orbach et al., 2000; Yoshida et al., 2009), highly conserved *AVRs* could have been selected to occupy euchromatic and repeat-poor regions. *AVR-Pi9*, a widely present effector among *M. oryzae* pathotypes, is located in a genomic region close to the chromosome 7 centromere, which is one of the most stable regions of the genome (Wu et al., 2015). Similarly, *AVR-Piz-t* is found at high frequency among pathotypes and located 230 kb from *AVR-Pi9* (Li et al., 2009). Effectors with highly variable presence/absence frequencies among pathotypes include *AVR-Pia*, *AVR-Pii*, *AVR-Pita* and *PWL* and are well-known to be located in unstable chromosomal regions such as subtelomeres prone to rearrangement (Kang et al., 1995; Orbach et al., 2000; Sweigard et al., 1995; Yasuda et al., 2006; Yoshida et al., 2009). Despite the broad spectrum in chromosomal locations, most of the effectors are expressed during the early stages of infection. AVR-Pi9 accumulates in the biotrophic interfacial complex structure and is translocated in the early stages of infection to the host cell (Wu 2015). A similar expression and translocation pattern was also observed for less conserved effectors such as AVR-Pia and AVR-Pita (Han et al., 2021; Khang et al., 2012; Sornkom et al., 2017).

TE activity is typically high in telomeric and subtelomeric regions (Croll & McDonald, 2012; Faino et al., 2016). Here, we show that *AVRs* with highly variable frequencies among pathotypes are indeed surrounded by TEs. The presence of TE can also affect epigenetic regulation through changes in heterochromatic structure (Aparicio et al., 1991; De Las Peñas et al., 2003; Fan et al., 2008). These results reinforce the idea that in *M. oryzae* different TE environments impact effector evolution in distinct ways. Our findings also indicate that certain TE superfamilies may have more affinity for specific *AVR*s loci, reflected also in the fact that the TE presence is largely conserved among pathotypes. This suggests that despite differences in TE activity among fungal plant pathogens (Gourlie et al., 2022; Oggenfuss et al., 2021; Shirke et al., 2016), including in *M. oryzae* (Lin et al., 2024; Nakamoto et al., 2023), the effector regions may have been selected for a variety of genomic niche features. The clearest evidence for TE insertions creating transitional states at effector loci was found for *AVR1-CO39*, with TEs likely contributing to the loss in MoO.

Overall, we show that *M. oryzae AVR* locus evolution was characterized by parallel and well differentiation dynamics in TEs. Spanning the spectrum of effectors exhibiting rapid changes in frequencies across *M. oryzae* pathotypes to conserved effector loci retained at high frequency and devoid of TEs. This highlights that host-mediated selection plays not only a role in *AVR* frequencies but that the genomic niche of the effectors displays likely similar associated dynamics. The dynamics of TE insertions within plant pathogens as a response to host selection remains understudied. Our work demonstrates though that TE dynamics can be an integral component of genomic niche evolution. In-depth tracking of effector niches will likely augment our ability to predict the durability of host resistance.

## Declarations

## Acknowledgements

We are grateful to group members for critical discussions.

## Data availability

The data analyzed in the frame of this study were retrieved from NCBI repositories as indicated in the Supplementary Tables S1 and S2. Draft genome assembly data is available from Zenodo (https://doi.org/10.5281/zenodo.15496085).

