## Supplementary Figure S1 for "Transposable elements create distinct genomic niches for effector evolution among *Magnaporthe oryzae* lineages"

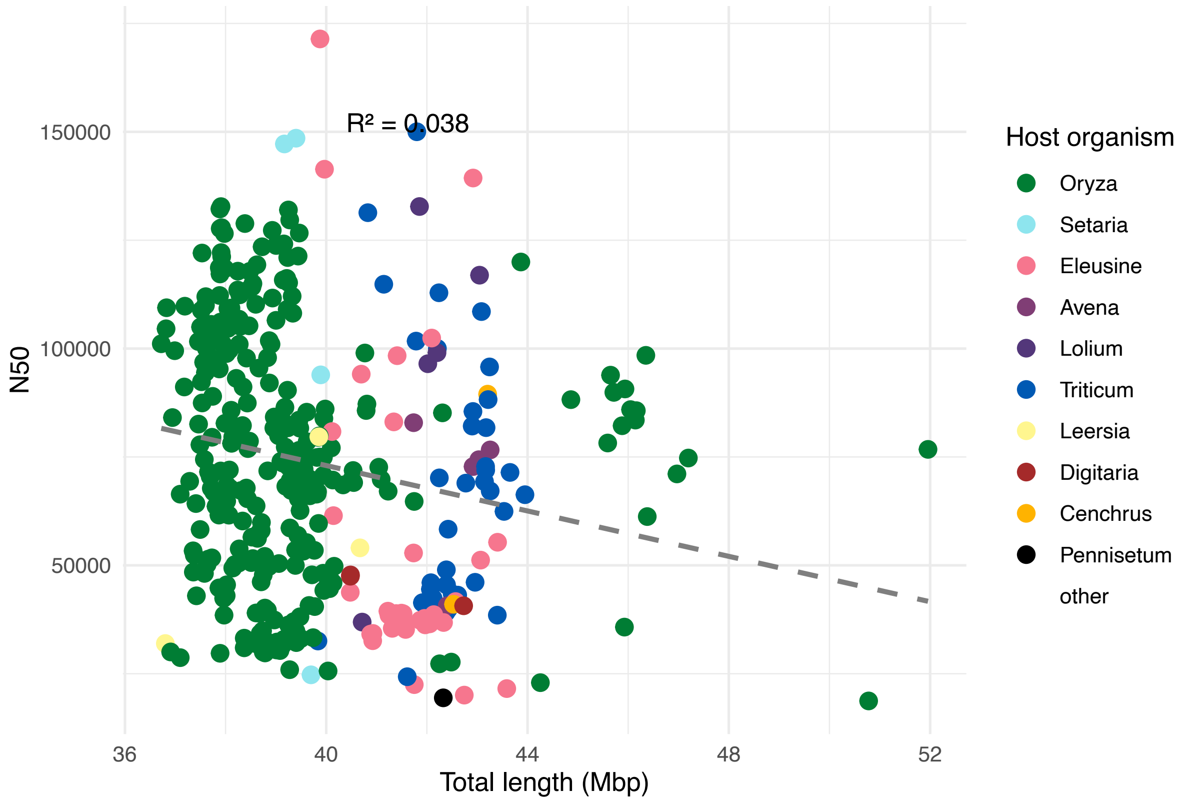


**Supplementary Figure S1:** Correlation between total genome assembly size and N50 metrics for *de novo* assembled genomes. Colors identify the reported host for each isolate.
